# Fundamental Limitations of Foundation Models in Single-Cell Transcriptomics

**DOI:** 10.1101/2025.06.26.661767

**Authors:** Srijan Atti, Shankar Subramaniam

## Abstract

Recent applications of foundation models in biology have focused on pretraining using large-scale single-cell datasets comprising millions of cells, across diverse patho-physiological states. These models are then fine-tuned for downstream tasks such as cell-type classification. In this study, we evaluated the performance of three widely-used foundation models in biology—scGPT, SCMAMBA-2, and Geneformer—and a statistical baseline (Seurat v5) on cell-type classification under Gaussian noise perturbation. We used two curated datasets, referred to as Myeloid (13k cells) and hPancreas (15k cells). Surprisingly, we found that the baseline performance of the foundation models was inferior to that of the statistical model, even without any added perturbation. Although we note that model size can affect performance, Geneformer’s accuracy outperformed scGPT and SCMAMBA-2 by 5% on average across all datasets despite having 40% fewer trainable parameters. Nonetheless, the statistical baseline still outperformed Geneformer by 9% in accuracy. Based on these findings, we hypothesized that the conventional training paradigm used by foundation models for single-cell tasks consistently underperforms statistical models due to a lack of essential biological context. To investigate this, we evaluated whether performance degradation stems from early data embedding steps—such as binning or gene normalization during tokenization, and sampling bias. First, to better understand tokenization-related artifacts, we introduced controlled Gaussian noise to gene expression values before tokenization, amplifying downstream distortions introduced by the tokenization process (all models were trained for identical step durations with identical hyperparameters). On the Myeloid dataset, following the introduction of Gaussian noise perturbation to 20% of cells, both scGPT and SCMAMBA-2 saw an 11% decrease in accuracy while Geneformer saw an 8% decrease in accuracy. This difference in performance may be due to the difference in encoding methods used by both models. scGPT and SCMAMBA-2 use a bin-based tokenization strategy, in contrast to Geneformer’s rank-value encoding, which normalizes gene expression values using a predetermined encoding constant. Although binning captures general trends in count data, it fails to preserve relative expression at the gene level, resulting in significant information loss. Applying the three models to the Myeloid dataset also revealed that scGPT’s prediction distribution is biased toward overrepresented cell types in the training data, while underrepresenting rarer classes. CD14 cells are overpredicted by 16% (among most abundant cell types) by scGPT. Geneformer, however, maintains a more stable prediction distribution with a 6% (CD14) increase and outperforms scGPT and SCMAMBA-2 by 26% in macro F1 score (unweighted metric). Based on our findings we assert that the gap in contextual encoding in bin-based tokenization is what contributes to the less-nuanced learning. Recent research that integrated cellular-ontology during training showed improved performance to both scGPT and Geneformer. Our results underscore a fundamental issue, that foundation models lack critical biological context that would allow for them to make the nuanced inferences required for complex biological analyses. The compression of single-cell data from raw counts to embedding vectors can span several orders of magnitude, and lead to significant loss of information. As a result, methods must adapt to prioritize contextual integration during tokenization to ensure sufficient information for the model.

## 1 Introduction

Foundation models, through their ability to develop rich contextual representations, have driven significant advancements across multiple domains —including language [1], medical imaging [2], time-series analysis [3], and genomics [4]. These models are typically trained on massive, heterogeneous datasets that are not limited to specific downstream tasks. Owing to the scale and scope of their training data, foundation models have demonstrated strong performance in diverse applications such as natural language processing and image generation [5]. As such, foundation models have led to substantial growth in the myriad of fields they have been applied to. These models have allowed for enhanced analysis in two key ways: first, by providing a framework capable of capturing fundamental features in pre-training data, they allow for robust fine-tuning for downstream tasks; and second, by using strong fundamental understanding to improve performance in low-data or noisy-data conditions.

More recently, foundation models have enabled significant advances in single-cell biology, particularly in tasks such as cell-type classification, gene interaction modeling, and dosage sensitivity prediction. [6]. Over the past five years, numerous foundation models have been developed for single-cell applications, including (but not limited to) scGPT [7], SCMAMBA-2 [8], Geneformer [9], scBERT [10], SCimilarity [11], and GeneCompass [12]. This underscores the critical role of foundation models in extracting meaningful insights from high-throughput omics data. Despite their promise, the effectiveness of these models rests on several critical assumptions: (1) that current tokenization strategies sufficiently preserve relevant biological information for effective pretraining; (2) that the datasets used for both pretraining and fine-tuning are large and diverse enough to support generalization; and (3) that the training objectives employed are well aligned with the biological complexity of single-cell transcriptomic data.

In this study, we evaluate whether existing single-cell foundation models are capable of developing meaningful biological reasoning. We focus on three representative models: SCMAMBA-2, scGPT, and Geneformer, two of which are based on transformer architectures, while the third employs a hybrid state-space model. We assess the quality of the learned embeddings on the widely used cell-type classification task. Additionally, we examine model robustness by benchmarking performance in low-data and high-noise regimes against a statistical baseline, Seurat v5.

## 2 Methods

### 2.1 Foundation Models Utilized

In this study we analyzed the performance of three models: scGPT [7], SCMAMBA-2 [8], and Geneformer [9] (Table 1). These models utilize different tokenization strategies including: (i) *bin-based*, in which each gene’s expression was discretized into a fixed number (50) of equal-width bins per cell; and (ii) *rank-based*, where genes were sorted by expression and encoded by their rank within each cell’s transcriptome. In the following sections, we provide a brief description of the foundation models, and the corresponding tokenization strategies employed.

**Table 1.** Key characteristics of the three foundation models.

| Model | Architecture | Tokenization | Pretraining Corpus |
| --- | --- | --- | --- |
| scGPT | 12-layer Transformer, 8 heads | Bin-based | ~33 M cells |
| SCMAMBA-2 | SSM (Mamba), linear scaling | Bin-based | ~57 M cells |
| Geneformer | 12-layer Transformer, 8 heads | Rank-based | ~30 M cells |

**scGPT** is a transformer-based model pre-trained on approximately 33 million non-cancer single-cell transcriptomes using a masked language modeling objective. The model comprises 12 encoder layers with 8 attention heads each and directly learns cell embeddings by conditioning masked-gene predictions on a predicted cell token. The model consists of approximately 51 million trainable parameters.

**SCMAMBA-2** employs a structured state-space model (SSM) backbone that, in essence, combines convolutional and recurrent computations to capture long-range dependencies with linear time complexity. Trained on a corpus of 57 million cells, SCMAMBA-2 contains approximately 60 million parameters and leverages the same bin-based tokenization as scGPT. To accommodate the inherently sequential SSM architecture, a bi-directional MAMBA block feeds the input sequence forward and reverse into two parallel SSM heads, whose outputs are averaged. This bi-directional system ensures that learned-parameters are position-agnostic to any input sequence.

**Geneformer** is a transformer-based foundation model pretrained on roughly 30 million non-malignant single-cell transcriptomes using a rank-based encoding scheme. Each input cell is represented as a ranked list of genes, where a gene’s position reflects its relative expression and is scaled by its corpus-wide expression distribution to emphasize cell-state–discriminative genes. The model uses 12 encoder layers with 8 attention heads and is trained via a masked gene prediction objective in which 15 % of tokens are masked at random. The model has roughly 40% fewer trainable parameters than scGPT, and roughly 50% fewer parameters than SCMAMBA-2.

All models were fine-tuned on an RTX6000 GPU for the same number of steps (depending on training set).

### 2.2 Tokenization Strategies

In the following sections, we describe the tokenization strategies used by the aforementioned foundation models.

#### 2.2.1 Bin-Based Tokenization

This tokenization approach is utilized by both scGPT and SCMAMBA-2 and has three components that are summed and returned as input for the model including gene token, gene expression bins, and condition tokens. As previously described [7][8], these components are calculated as follows. **Gene Token:** To provide the model with information on the identity of individual genes, each gene is assigned a unique integer identifier (gene token). These gene tokens are selected from a predefined vocabulary constructed from the union of gene sets observed in the pre-training dataset. This approach ensures consistency across datasets while accommodating variation in gene coverage due to differences in sequencing technologies or processing pipelines. **Gene Expression Binning:** To incorporate the magnitude of the expression while mitigating batch-specific variation, the raw gene expression counts are first logarithmically transformed, then rank-ordered within each cell, and discretized into bins *b*. This binning procedure normalizes the relative expression levels of genes, allowing the model to interpret the expression in a context-invariant manner. As a result, the model captures the relative importance of each gene within a cell, regardless of the depth of the sequencing or some level of technical noise. **Condition Token:** To provide extraneous information to the model beyond gene identity and expression, condition tokens are introduced. These tokens are position-wise embeddings that encode meta-information associated with each gene. These tokens capture experimental or functional annotations, such as pathway membership, perturbation status, or other contextual variables relevant to the modeling task. Each condition is assigned a unique integer identifier, allowing heterogeneous biological conditions to be represented uniformly across diverse datasets. By aligning condition tokens positionally with gene tokens, scGPT enriches its representation space with biologically meaningful priors that guide learning across multicontext settings.

For each gene position *j* in a given cell, the final input embedding *e*_*j*_ is computed as the sum of the three distinct embedding vectors:

*E*_*gene*_(*g*_*j*_): the embedding of *g*_*j*_, the gene token.

*E*_*expr*_(*x*_*j*_): the embedding of the binned expression value *x*_*j*_, representing the gene’s relative expression level within the cell.

*E*_*cond*_(*c*_*j*_): the embedding of the condition token *c*_*j*_, which incorporates contextual metadata such as perturbation labels or pathway annotations.

These components are combined with element-wise addition:

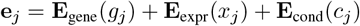

The resulting sequence {*e*_1_, *e*_2_, …, *e*_*M*_} serves as the input to the foundation model’s encoder, where M denotes the number of selected genes per cell.

#### 2.2.2 Rank Value Encoding

In contrast to the bin-based tokenization method that scGPT and SCMAMBA-2 use, Geneformer utilizes a rank value encoding scheme that aims to capture relative gene expression information. This rank value encoding method generates encoding vectors as follows. **Global Gene Normalization Factors:** The encoding method aims to capture the relative importance of genes within each cell, normalized by global expression patterns from the large pretraining corpus. This global gene normalization factor, which we denote 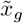 is calculated over, in this case, the Genecorpus-30M dataset. For each gene *g*, we calculate its non-zero median expression across all cells in which it is detected:

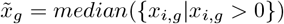

where *x*_*i,g*_ denotes the raw trasncript count of gene *g* for cell *i*. Zeros are excluded from the distribution to prevent any bias as a result of tissue or condition-specific dropout effects. **Per-Cell Normalization and Ranking:** For each new cell *i* encountered following pre-training, the gene expression is normalized by the total transcript count in that cell to correct for sequencing depth. Following this, each gene’s expression 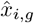 is divided by 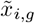 The rank of each gene (*r*_*i,g*_) based on the computed *x*^*′*^_*i,g*_ is computed. This method allows for the prioritization of relatively over-\expressed genes. **Tokenization and Sequence Construction:** Each gene is finally assigned a unique integer token id (*g*) from a fixed vocabulary of genes that are based on the pre-training corpus. For each cell, genes are ordered by *r*_*i,g*_ and their corresponding tokens are arranged into a sequence 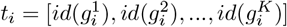 where 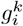 denotes the *k*-th highest normalized expression in cell *i*, and *K* is the number of expressed genes in that cell.

### 2.3 Datasets Utilized for Fine-Tuning

To evaluate the performance of each model on the task of cell-type classification, we used three datasets that have been previously utilized for model evaluation [7] [8]. These datasets consist of cells from human pancreas (hPancreas) [13] and a tumor infiltrating Myeloid (Myeloid) [14] dataset (Table 2).

**Table 2.** Training and test set sizes for the three datasets used in this study.

| Dataset | Train Size | Test Size |
| --- | --- | --- |
| hPancreas | 10,600 | 4,218 |
| Myeloid | 9,748 | 3,430 |

All input data were subjected to standard single-cell RNA sequence preprocessing, including normalization (log normalization), correction for confounders, and selection of highly variable genes features prior to tokenization.

### 2.4 Gaussian Noise Perturbation Experimental Procedure

To evaluate the robustness and generalization ability of each foundation model, we performed test-time Gaussian noise perturbation. This experiment was designed to identify the effect that tokenization had on model performance with unstable data.

To simulate increasing levels of technical noise in the test data, we added Gaussian noise to the input expression profiles at inference time. The noise, which has a standard deviation of the median count of the nonzero elements in the dataset, is applied to *n*% of the dataset in 10% increments (i.e. *n* = 10%, 20%, …, 100%). During this test-time, model weights remain fixed—having been fine-tuned on the original noise-free training set. This setup evaluated model robustness to perturbations not seen during training, specifically amplifying post-tokenization artifacts by introducing pre-tokenization noise.

At *n* = 30%, we observed a breakdown in cluster integrity in the PCA space, with underrepresented cell types becoming indistinct or merging with dominant populations. This also marked the point of failure for our statistical baseline and motivated its designation as a critical threshold.

This study primarily evaluated the efficacy of each model to perform cell-classification using accuracy, precision, recall, and F1 scores. Due to class imbalances in the data, we utilize macro-weighting, calculated as: 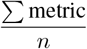), for all metrics except accuracy. Macro weighting is beneficial because it is a class-agnostic, or unweighted, method for metric computation.

## 3 Results

### 3.1 A Comparison of Fine-Tuning Classification Performance

We evaluated the performance and learning capabilities of three foundation models: scGPT, SCMAMBA-2, and Geneformer for the task of cell-type annotation (see Methods) on two datasets, referred to as hPancreas (14.8k cells) and Myeloid (13.2k cells). We first extracted embeddings generated by the pre-trained foundation model and passed them into functionally identical classification heads. We compared their performance with an often utilized and well-established statistical model, Seurat v5.

The results of the learned classification evaluated on their accuracy, macro precision, macro recall, and macro F1 scores (Table 3, Figure 1). We found that all three foundation models performed significantly poorer than Seurat on all datasets. We noted that Geneformer performed approximately 5% better than both SCMAMBA-2 and scGPT on average across both datasets, for accuracy. This is in spite of having nearly 40% less trainable parameters. The statistical baseline (Seurat v5) performed consistently better than all models for all performance metrics. Specifically, Seurat outperformed Geneformer by a margin of ~ 15% in accuracy.

**Table 3.**
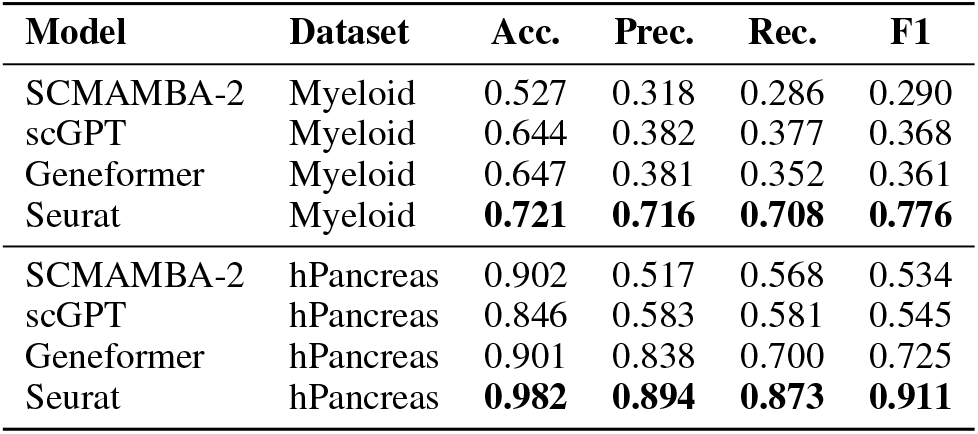
Performance metrics for each model on the Myeloid and hPancreas datasets. Best metrics highlighted.

**Figure 1.**
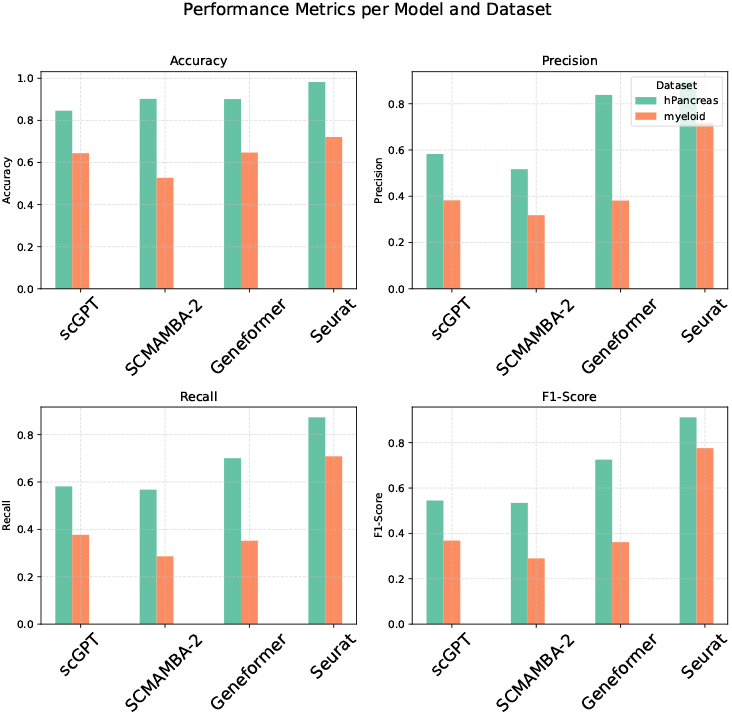
Model performance across all metrics and datasets.

### 3.2 Evaluating the Effects of Noise Perturbation

To elucidate the underlying reasons for the poor training performance observed, we assessed the robustness of features learned by each foundation model. Specifically, we introduced Gaussian noise perturbation to pre-tokenization data to amplify post-tokenization artifacts, and analyzed the differences between each model’s tokenization method and its effect on performance. We plotted the response to Gaussian noise perturbation for the same four key metrics: accuracy, macro precision, macro recall, and macro-F1 score against the *n*% of counts to which Gaussian noise was added (Figure 2).

**Figure 2.**
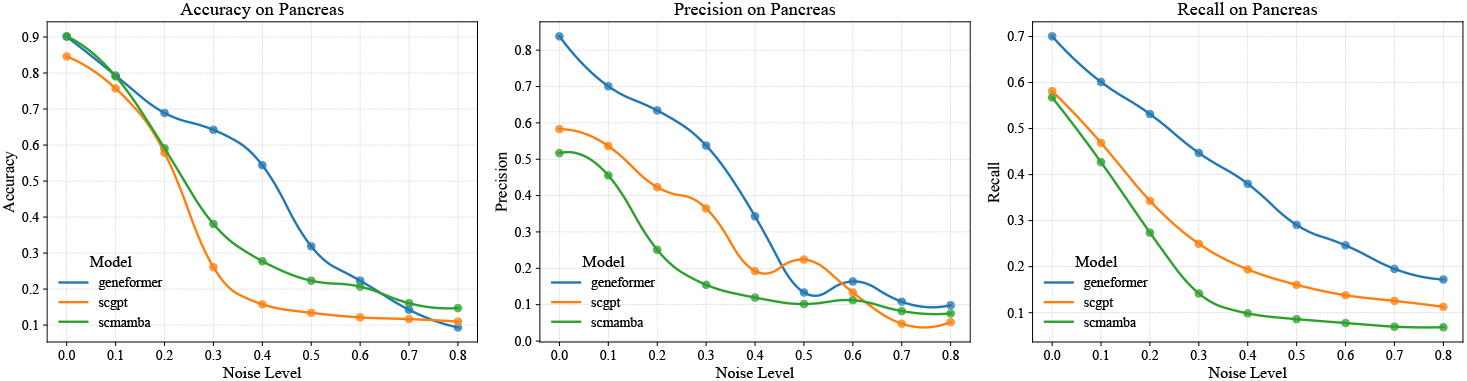
Plot of performance metric vs. *n*% of test dataset modified by Gaussian noise. *n*=[0–80]%

Our analysis, following the introduction of Gaussian noise perturbation, showed that Geneformer, despite its smaller size, still performed better on the hPancreas and Myeloid datasets. We observed an approximately 59% degradation in accuracy for scGPT’s predictions following the introduction of 30% Gaussian noise. Comparatively, we see a reduced effect for Geneformer’s predictions, with a lesser ~ 25% decrease in accuracy for the hPancreas dataset.

We also applied Gaussian perturbation to the Myeloid dataset and observed that Geneformer performed consistently better (Figure 3). Geneformer observed an ~25% decrease in performance following the introduction of 30% Gaussian noise, while scGPT observed an ~35% decrease in accuracy.

**Figure 3.**
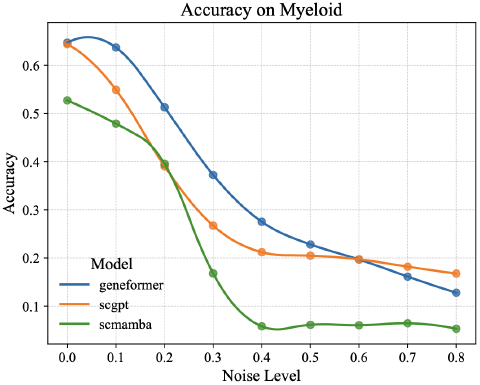
Accuracy vs. percent Gaussian noise (*n*=[0–80]%).

**Figure 4.**
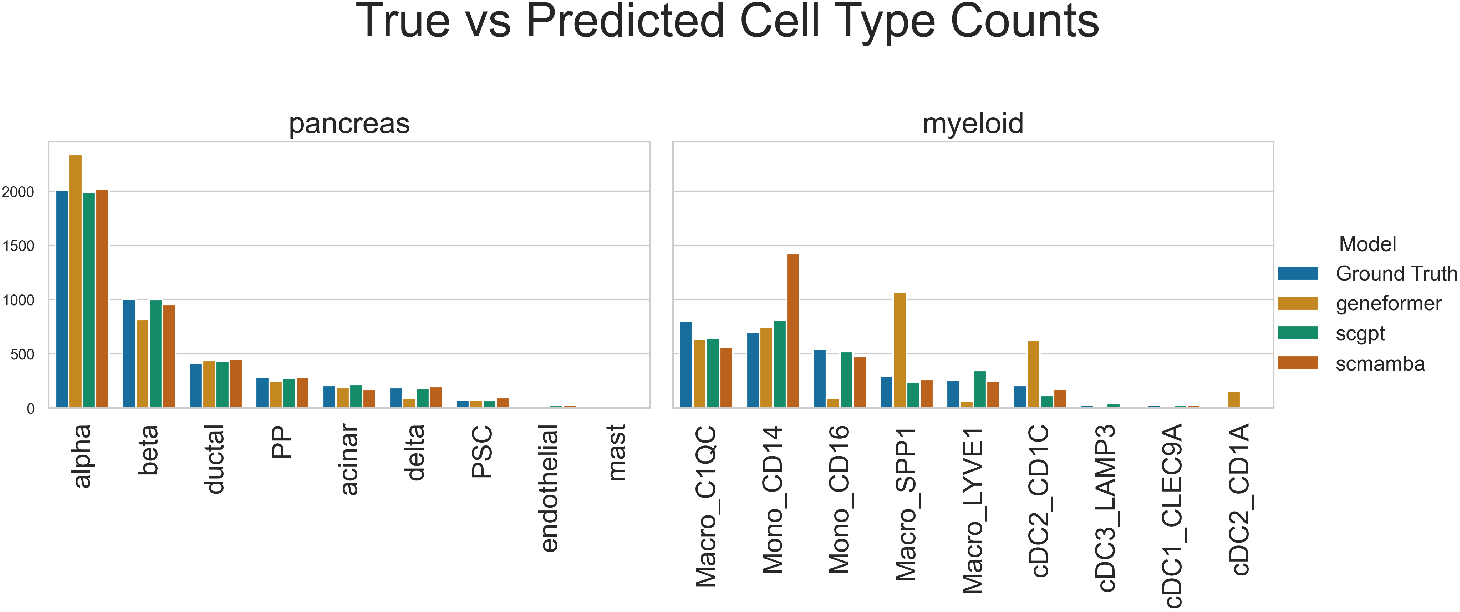
Distribution of predicted counts for each class type vs. true counts for each class type.

### 3.3 Effects of Sampling Bias During Fine-Tuning

The raw input data that was passed into the foundation models during fine-tuning was heavily class-imbalanced, and thus it was important to assess the impact of sampling bias on fine-tuning of single-cell foundation models.

We observed that the classification distributions for all models on the hPancreas dataset were quite stable with no major fluctuations present. However, the Myeloid dataset yielded different results. In particular, SCMAMBA-2 predicted CD14+ Monocyte at nearly double the rate of true CD14 classification. CD14+ Monocytes are one of the most abundant class in the Myeloid dataset, indicating that SCMAMBA-2 likely overfit to the fine-tuning dataset. scGPT and Geneformer also saw increased prediction of CD14+ Monocytes at a more modest rate of ~16% and ~6%, respectively. Additionally, Geneformer severely over predicted Macrophage SPP1, indicating training uncertainty caused by underrepresentation of the SPP1 cell type in the training dataset.

Although the distribution of Geneformer’s predictions for the Myeloid dataset yielded more volatile results, its macro metrics remained on-par with scGPT (see Table 3). Notable was that Geneformer performed significantly better on macro metrics (precision, recall, and F1). Geneformer —on the hPancreas dataset in specific —performed 10-20% better than scGPT for all macro metrics indicating that, especially for underrepresented classes, Geneformer did a better job at classification. Thus, we conclude that, foundation models, regardless of tokenization method, are not resistant to train-set sampling bias during fine-tuning. This sampling bias leads to sub-par performance and skewed prediction distributions, particularly towards overrepresented classes.

## 4 Discussions

In this study, we demonstrate that the performance of foundation models—specifically scGPT, SCMAMBA-2, and Geneformer—is currently inferior to that of conventional statistical methods such as Seurat v5. Fine-tuning these models across three datasets (Myeloid and hPancreas) revealed a consistent and notable performance gap between statistical approaches and the tested foundation models. Furthermore, our preliminary analysis uncovered performance differences among the foundation models themselves. To investigate the source of this disparity, we introduced pre-tokenization noise to simulate post-tokenization artifacts, thereby probing the resilience of each model’s encoding strategy. When presented with these perturbed inputs, Geneformer demonstrated markedly greater robustness compared to scGPT and SCMAMBA-2. Notably, Geneformer achieved this despite having approximately 40% fewer trainable parameters than the other two models. We attribute this robustness to Geneformer’s rank-value encoding strategy, which incorporates a global gene normalization factor. This design enables the model to assess gene expression in the context of its median value within the training dataset, thereby preserving biologically meaningful relationships such as gene upregulation or downregulation. In contrast, the bin-based encoding methods used by scGPT and SCMAMBA-2 reduce expression levels into 50 equidistant bins following normalization. While this strategy can buffer against noise through discretization, it risks oversimplifying cell states and obscuring subtle biological signals. Our findings emphasize the importance of integrating biologically relevant context into model design. Recent studies have demonstrated that incorporating cell ontological information during model encoding significantly improves performance over approaches like Geneformer and scGPT [15]. These insights collectively underscore a crucial next step for advancing machine learning-driven single-cell analyses: the development of encoding strategies that faithfully preserve and leverage the inherent biological complexity of cellular systems.

